# Collective responses of flocking sheep to a herding dog

**DOI:** 10.1101/2024.05.24.595762

**Authors:** Vivek Jadhav, Roberto Pasqua, Christophe Zanon, Matthieu Roy, Gilles Tredan, Richard Bon, Vishwesha Guttal, Guy Theraulaz

## Abstract

Across taxa, group-living organisms exhibit collective escape responses to stimuli varying from mild stress to predatory pressures. How exactly does information flow among group members leading to a collective escape remains an open question. Here we study the collective responses of a flock of sheep to a shepherd dog in a driving task between well-defined target points. We collected highresolution spatio-temporal data from 14 sheep and the dog, using Ultra Wide Band tags attached to each individual. Through the time delay analysis of velocity correlations, we identify a hierarchy among sheep in terms of directional influence. Notably, the average spatial position of a sheep along the front-back axis of the group’s velocity strongly correlates with its impact on the collective movement. Our findings demonstrate that, counter-intuitively, directional information on shorter time scales propagates from the front of the group towards the rear, and that the dog exhibits adaptive movement adjustments in response to the flock’s dynamics. Furthermore, we show that a simple shepherding model can capture key features of the collective response of the sheep flocks. In conclusion, our study reveals novel insights on how directional information propagates in escaping animal groups.

## I. INTRODUCTION

The ability of groups of organisms to detect threats or predator attacks and then coordinate their collective movements to escape is observed in many species living in groups such as swarms of insects [1], schools of fish [2– 5], flocks of birds [6–10], and herds of mammals [11– 13]. These patterns of collective motion often confuse the predator and increase the survival of the prey [14– 18].

These properties, sometimes referred as a form of collective intelligence [19, 20], emerge at the group level from specific behavioral interactions between individuals [21– 29]. Studying these interactions can provide insight into the way information spreads across groups of organisms in response to predator attacks and into the mechanisms by which animals coordinate their actions [30–35]. This knowledge is also fundamental to understanding how selective pressure acting at the individual level promotes flocking behaviors [14, 36–40].

Studying collective escape phenomena in the field is a challenging task. Not only predator attacks are unpredictable and rare events, but simultaneously obtaining data on the trajectories of the predator and individuals within the group during these attacks can be extremely complex. To overcome these difficulties, some studies simulate threats, via various methods such as a human approaching antelopes [13] or robotic models of predators approaching a flock of birds [41] or a school of fish [42], and instigate collective response.

Collective responses can be investigated via the novel system of sheep herding, where interactions of a flock of sheep with a herding dog offers the opportunity to study the impact of a controlled threat on the collective movements of an animal group [43]. As a matter of fact, herding dogs have retained only some sequences of the predation behavior of wild canids. They display motor behavioral patterns such as searching and orienting, fixing, following, approaching, and finally chasing that are observed in related wild canids such as wolves and coyotes. But unlike the complete sequence characteristic of these species, the capture and killing of prey are absent in herding dogs [44]. Moreover, several studies have reported that in the presence of a dog, sheep exhibit fearful behavior [45, 46] and their level of stress increases alongside an increase in plasma cortisol concentration [47], suggesting that sheep perceive a herding dog as a threat. Since the behavior of the dog is controlled by the shepherd, it is thus possible to carry out a large number of replications of the same experimental situation.

Here, we investigate the collective responses of a flock of sheep to the behavior of a herding dog. As the dog gets closer to the flock, it induces an avoidance reaction from the sheep. We analyze how this behavioral reaction to the perceived threat propagates within the flock and affects the collective movements of sheep. In our experiments, a trained border collie was responding to the verbal commands of the shepherd and used to guide a flock of sheep (*N* = 14) from an initial location to a target location. We collected the positions and orientation of all sheep, the dog and the shepherd with a Ultra wide band-based real-time location system during several dozen of trips performed by the flock. We first analyze the collective movements of the flock when interacting with the dog. We then analyze the directional correlation between the dog and the flock, as well as among the individual sheep in order to determine how information propagates within the flock. We finally used a modified version of a shepherding model to show that simple interaction rules between sheep and between sheep and dog can reproduce key features of collective escape response and information flow observed in our data.

## II. MATERIALS AND METHODS

### Study species

14 female merino sheep (*Ovis aries*) aged 4 years and with a mean body weight of 75 kg and mean body length of 1.2 m were used in the experiments.

We also used a 8 years old border collie specially trained to herd sheep and whose behavior during the experiments was managed by a professional handler (Fig. 1**a**).

**FIG. 1.**
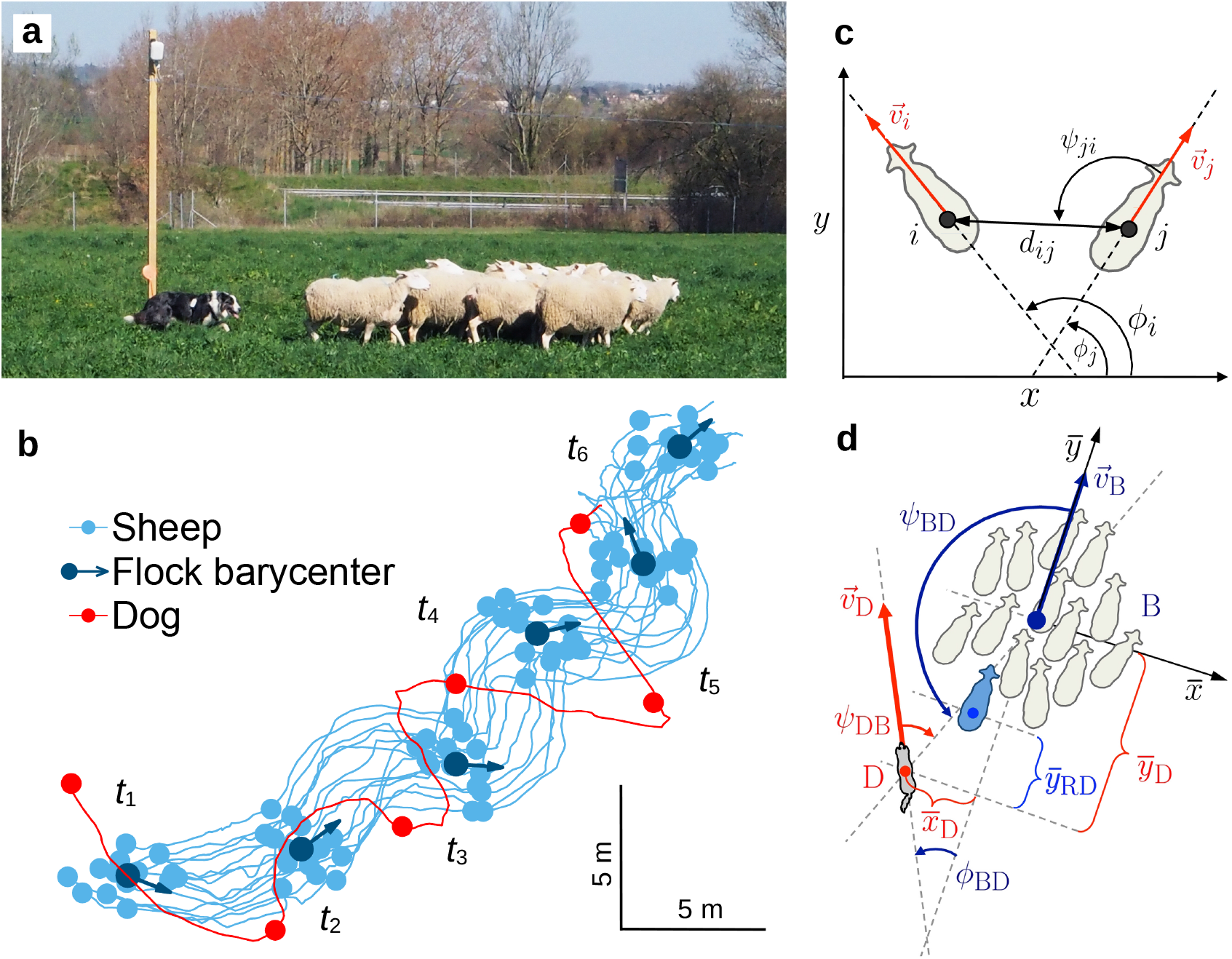
(**a**) Flock of 14 females merino sheep (*Ovis aries*) and sheepdog (*Border collie*), equipped with Ubisense tags to track their positions while moving in a field of 100 m×50 m, passing in front of a pole on which a Ubisense Ultra Wide Band sensor is fixed. (**b**) Trajectories (solid lines) of the 14 sheep (light blue) and the dog (red) along 26 seconds of a driving event, with their positions (circles) shown each 5 s and labeled from *t*_1_ to *t*_6_. The barycenter of the flock and its instantaneous direction of motion are shown in dark blue. (**c**) Velocity vectors 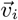 of two sheep *i* and *j*, with their heading angles ϕ_*i*_ and ϕ_*j*_, the angle with which sheep *j* perceives sheep *i*, and the distance between them *d*_*ij*_ . (**d**) Velocity vector of barycenter of the flock 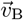, velocity vector of the dog 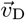, heading difference between both vectors ϕ_BD_, angle the barycenter has to turn to perceive the dog *ψ*_BD_, position of the dog 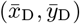 in the system of reference of the barycenter and in the direction of motion of the flock (given by 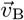), and distance of the rear sheep of the flock (sheep in blue color) to the dog, in the direction of motion of the flock, 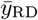.

### Tracking system

We used a real-time location system developed by Ubisense based on Ultra-Wide Band (UWB) signals triangulation [48] to track the movement trajectories of each individual sheep, the dog and the shepherd. Each sheep was equipped with two tags attached by a clip to the fleece along the axis of the vertebral column, which made it possible to get both the position and the orientation of the animal. The dog and his handler were also equipped with four and two tags respectively (Supplementary Fig.1). Ubisense tags are miniaturized circuits powered with batteries that operate in 6–8 GHz frequency band for localization and emit UWB wave trains that are then received and processed by a set of sensors. In our experiments, the tracking system included 8 sensors uniformly distributed around the experimental pasture and fixed 3 m from the ground. The sensors are UWB signal receivers connected through low-latency Ethernet cables to a server that processes sensor data into tags position. Each tag is localized at 2Hz with a typical error of less than 30cm.

### Herding events

Experiments were performed on a flat rectangular pasture of 100 m length and 50 m width. The task consisted for the shepherd and his dog to *drive* the flock between two locations A and B, 60 m apart. We performed a total of 26 round trips during which the flock was taken from place A to place B and then brought back to place A. Each round trip lasted 2 minutes on average. The controlled movement phases of the flock were interspersed with five rest periods of 30 to 60 minutes each to allow the dog to recover and maintain sufficient motivation to carry out the task. We focus our analysis on the collective movements during which the dog was driving the sheep and staying behind the flock relative to its direction of movement. While staying in this position, it may happen that the dog regularly alternates its direction of movement from left to right and vice versa following the shepherd’s orders.

### Ethics

Experiments were approved by the local Ethics Committee for Animal Science and Health and were performed at the experimental domain of Langlade (INRA, UMR 1388 GenPhySE), Pompertuzat, France, under permit APAFiS SSA 2017 005 in agreement with the French legislation.

## III. RESULTS

### A. Collective behavior of sheep and their reaction to the dog

Fig. 1**b** shows the trajectories of the *N* = 14 sheep (blue lines) and the dog (red) over a 26-seconds herding event, and the instantaneous positions of each individual at 6 equispaced instants of time *t*_1_, …, *t*_6_. During this particular event, the sheep move away from the dhog while remaining quite cohesive, with the dog staying behind the flock and exhibiting a zigzag motion (see also Supplementary video S1 and Supplementary video S2).

We characterize the sheep collective behavior by means of three observables: the group cohesion *C*, given by the mean radius of the group, the polarization *P*, which measures the degree of alignment of the individual sheep, and the elongation *E*, which measures the ratio length/width of the sheep flock with respect to its direction of motion (see Supplementary Fig.2). The position of the dog with respect to the sheep flock is described by the relative position of the dog with respect to the barycenter of the flock in the direction of motion of the flock, and the distance of the dog to the rear sheep in the flock in the direction of motion of the group.

We denote by 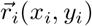 and 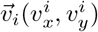 the position and velocity vectors of individuals, respectively, with *i* = 1, …, *N* for the sheep, *i* = D for the dog and *i* = B for the barycenter of the sheep flock given by 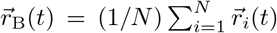. We consider that the heading of an individual is given by its direction of motion, *i*.*e*., by the angle of its velocity vector 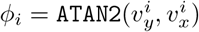 (Figs. 1**c, d**). Then, we define

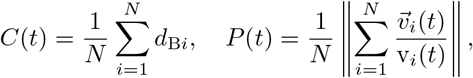

where 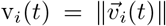 is the speed of individual *i* and 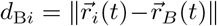 is the distance between the barycenter and *i*.

The viewing angle of *i* defined as the angle with which *i* perceives *j*, quantifies how much the velocity vector 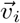 has to turn to point towards *j, ψ*_*ij*_ = θ_*ij*_ − ϕ_*i*_, where θ_*ij*_ = A*T*AN2(*y*_*j*_ *y*_*i*_, *x*_*j*_ *x*_*i*_) is the angle that the vector going from *i* to *j* forms with the horizontal x-axis. The alignment of two individuals is measured by the difference of their headings, ϕ_*ij*_ = ϕ_*j*_ ϕ_*i*_. Positive angles are measured counterclockwise, and negative angles are measured clockwise.

The average behavior of the flock is described by the velocity vector of the barycenter 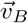 (Fig. 1**b**, dark blue arrow). This vector defines the ordinate axis of an oriented system of reference centered on B (Fig. 1**d**). Denoting the variables in this system of reference with a bar, we have, for *i* = 1, …, *N*, D:

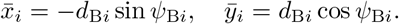

The length and the width of the group in its direction of motion can then be easily expressed using these variables, so that the elongation can be defined as

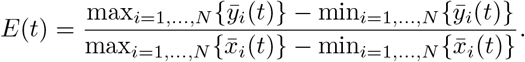

The sign of 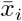 reveals the side where individual *i* is located with respect to the direction of motion of the sheep flock, and 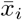 shows how far the individual is from this direction. In particular, 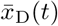 reflects the lateral oscillations of the dog with respect to the direction of motion of the flock. Along this direction, the distance of the dog to the barycenter of the flock is given by 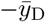, and the distance of the dog to the rear sheep of the flock by 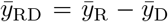, where *R* refers to the rear sheep in this direction, that is, 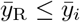 for all *i*. Negative values of 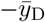 and 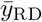 mean respectively that the dog is in front of the barycenter or in front of the rear sheep.

We are first interested in describing how the sheep responds to the presence and the movements of the dog. During the experiments, sheep were not continuously exposed to the dog and spent most of the time at rest in a more or less random spatial configuration. In turn, in presence of the dog, the sheep move and align to each other. Fig. 2 shows that two phases can be identified in the whole set of data captured during the experiments: an active phase (upper dark region) where both speed and alignment are high, and a passive phase (bottomleft dark region), where both speed and alignment are small. Supplementary Fig.3 also shows that speed and alignment are positively correlated. We therefore focus our analysis on the active phase.

**FIG. 2.**
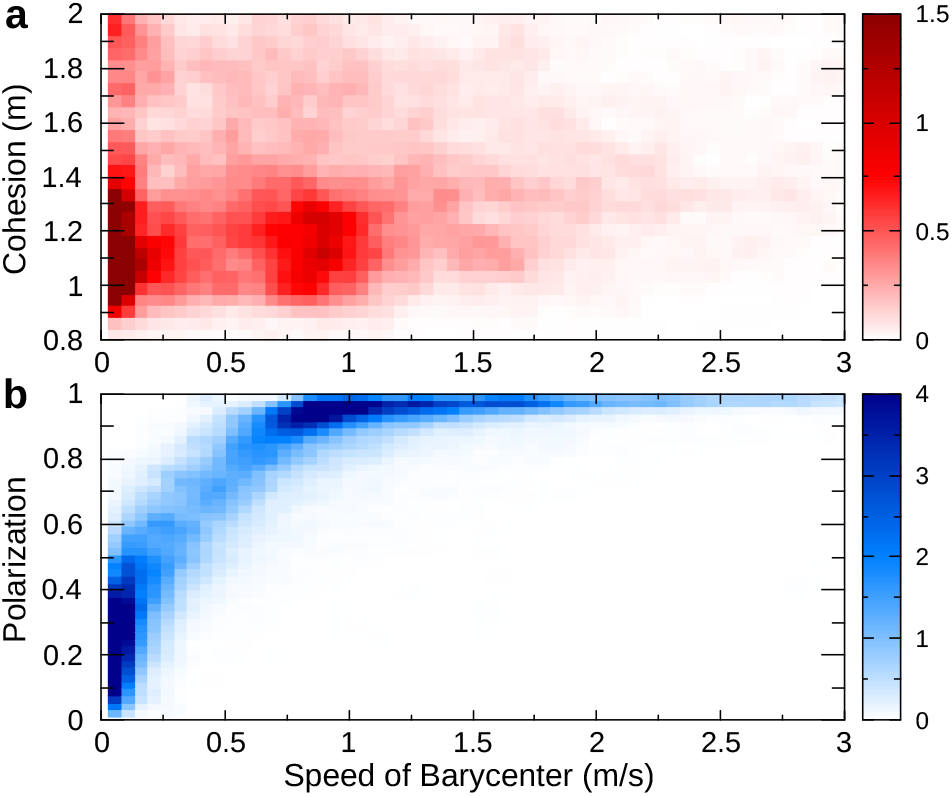
(**a**) Cohesion and (**b**) Polarization of the flock as a function of the speed of its barycenter v_B_. Two phases can be distinguished: an active phase at high speed and high polarization (v_B_ ≈ 1ms^−1^, *P* ≈ 0.9), and an inactive phase in which sheep are at rest and not aligned (v_B_ ⪅ 0.3ms^−1^, *P* ≈ 0.1–0.4). This figure includes all the data recorded during the experiments. The data used to analyse the sheep response to the dog correspond to the active phase.

Fig. 3**a** shows that, during the sequence depicted in Fig. 1**b**, sheep speed is about 1.3 ms^−1^, smaller than the dog mean speed, 2 ms^−1^. Sheep remain highly cohesive during the drive, *C* ≈ 1.2 m, which corresponds the typical body length of sheep (Fig. 3**b**); they remain well aligned, with *P*(*t*) > 0.6 all the time (Fig. 3**c**), which is substantially above the expected polarization value of *N* non-interacting individuals, *P*_0_ ≈ 1/ *N* = 0.27 ([49]). Flock adopt an oblong shape perpendicular to the direction of motion, with *E*(*t*) smaller than 1 most of the time (Fig. 3**b**).

**FIG. 3.**
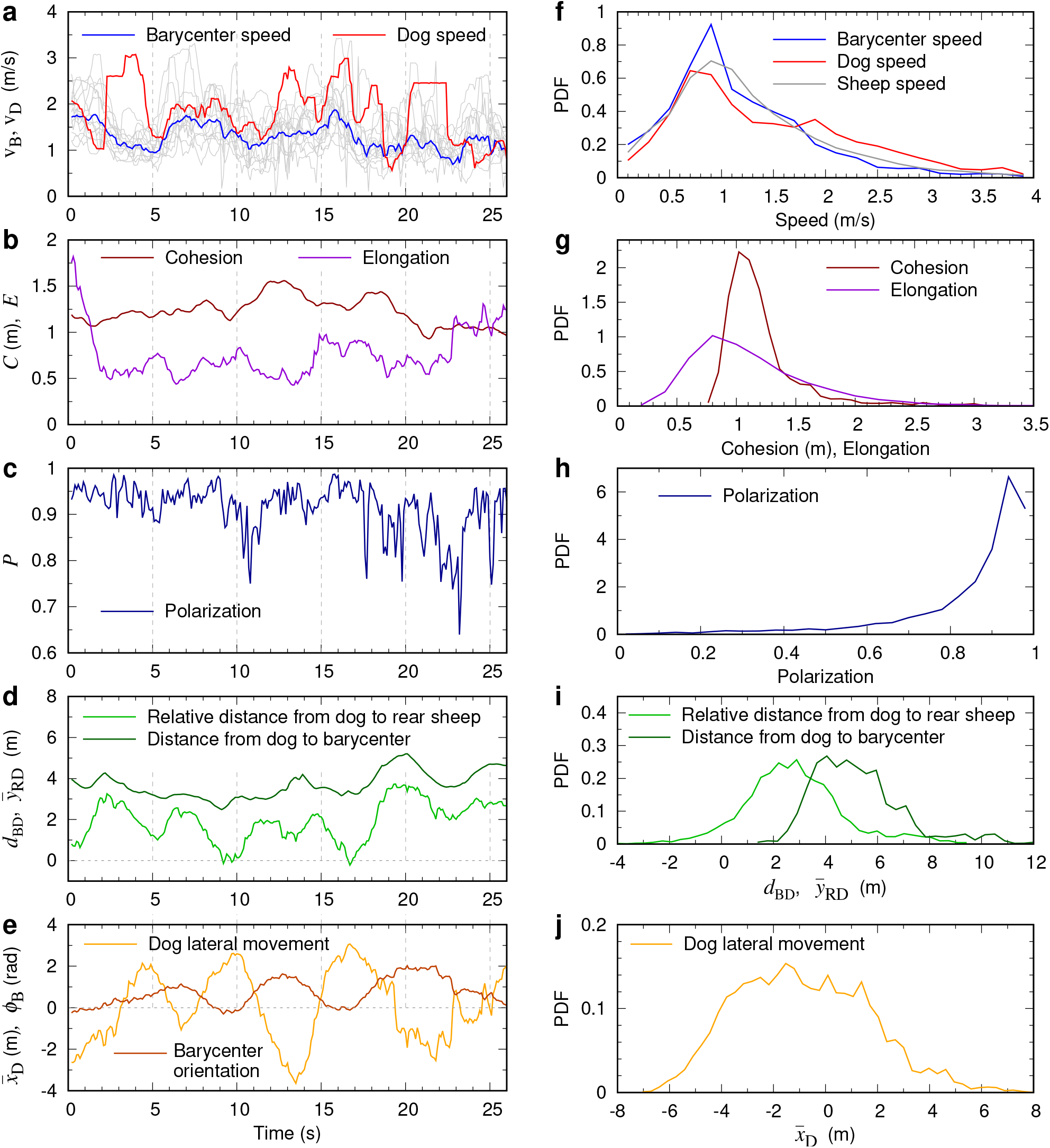
Time series (**a-e**) and probability density functions (**f-j**) of the observables characterizing collective behavior of sheep and their reaction to the dog: (**a**) Speed of the barycenter of the flock v_B_(*t*) (light blue), dog v_D_(*t*) (magenta), and individual sheep (gray lines). (**b**) Cohesion *C*(*t*) (red) and elongation *E*(*t*) (purple). (**c**) Polarization *P* (*t*) (blue). (**d**) Relative distance 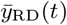 from the dog to rear sheep in the direction of motion of the flock, given by velocity vector of the barycenter (green), and distance *d*_BD_(*t*) from barycenter to the dog (dark-green). (**e**) Lateral movement of the dog with respect to the direction of motion of the flock 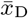 (orange), and orientation of the barycenter ϕ_B_(*t*). Vertical gray dashed lines show the instants of time shown in Fig. 1b. (**f-j)** Probability density function (PDF) corresponding to the observables shown in the left column. Mean ± SD values: ⟨*C*⟩ = 1.21 ± 0.34 m (∼body length); ⟨*P*⟩ = 0.85 ± 0.17; ⟨v_B_⟩ = 1.3 ± 0.002 ms^−1^; ⟨v_D_⟩ = 1.5 ± 0.01 ms^−1^; Mode of *E* = 0.8.

The dog remains always behind the flock barycenter and almost always behind every sheep with respect to the direction of motion of the flock, i.e. 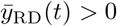 most of the time (Fig. 3**d**), and at a distance to the barycenter (*d*_DB_(*t*)) that varies ≈ 3–5 m (Fig. 3**d**). The dog exhibits wide zigzags with respect to the field (Fig. 1**b**), which are lateral movements with respect to the direction of motion of the flock, with 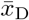 displaying wide oscillations between negative (dog is at the left side of flock direction of motion) and positive (dog is at the right side) values, with an amplitude often larger than 2 m (Fig. 3**e**). These oscillations are also visible in the trajectory of the flock, as shown by the changes of variation of its direction of motion given by the barycenter heading ϕ_B_ (Fig. 3**e**).

In order to know the extent to which these behavioral features can be generalized, we have measured the probability density functions (PDF) of the observables defined above (Fig. 3). Indeed, the two PDF of sheep speed and dog speed are quite similar, although the dog often reaches speed values that are quite larger than those of the sheep (see the bump at 2 ms^−1^ in the red curve in Fig. 3**f**). However, dog speed is more often smaller than sheep speed, with a peak of v_*D*_ located at 0.8 ms^−1^, below the location of the peak of v_*B*_ at 0.9 ms^−1^. In turn, the dog often reaches a speed higher than 2 ms^−1^, yielding a mean speed of 1.5 ms^−1^, slightly higher than the mean speed of sheep, 1.3 ms^−1^ (Fig. 3**f**).

The flock remains cohesive, with a mean radius ⟨*C*⟩ = 1.21 m (~body length), SD = 0.34 m, with a peak in elongation at *E* = 0.8 (Fig. 3**g**), and highly polarized, ⟨*P*⟩ = 0.85, SD = 0.17, as we selected the active phase for our analysis (Fig. 3**h**). The distance between the flock’s barycenter and the dog varies over a wide range, from *d*_BD_ ≈ 3 to 7 m (Fig. 3**i**). The dog is well behind the flock in the direction of motion of the flock, typically at a distance between 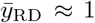 and 4 m from the rear sheep (Fig. 3**i**), and only in a small number of cases the dog surpasses the rear sheep 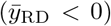. The wide lateral oscillations of the dog have a typical amplitude similar to the one observed in the illustrative sequence shown in Fig. 1**b**, from − 4 to 2 m, and can be even more pronounced, with 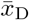 ranging from − 6 to 6 m (Fig. 3**j**). Note that the negative/positive asymmetry is simply due to the limited number of experiments, as no left/right asymmetry is expected in sheep or dog behavior.

Fig. 4 shows the impact of these wide lateral oscillations in the PDF of the angles with which the barycenter and the dog perceive each other. The PDF of the angle between the direction of the barycenter and the dog is wide, with values of |*ψ*_BD_| greater than 120^∘^ bing quite frequent, implying that the dog is behind the flock; conversely, the flock is ahead of the dog most of the times, although the distribution is relatively wide (Fig. 4**a**). Similarly, the PDF of the alignment between the barycenter and the dog headings is relatively wide, having practically the same shape than *ψ*_DB_, showing frequent misalignments larger than |ϕ_BD_| = 45^∘^ (Fig. 4**b**).

**FIG. 4.**
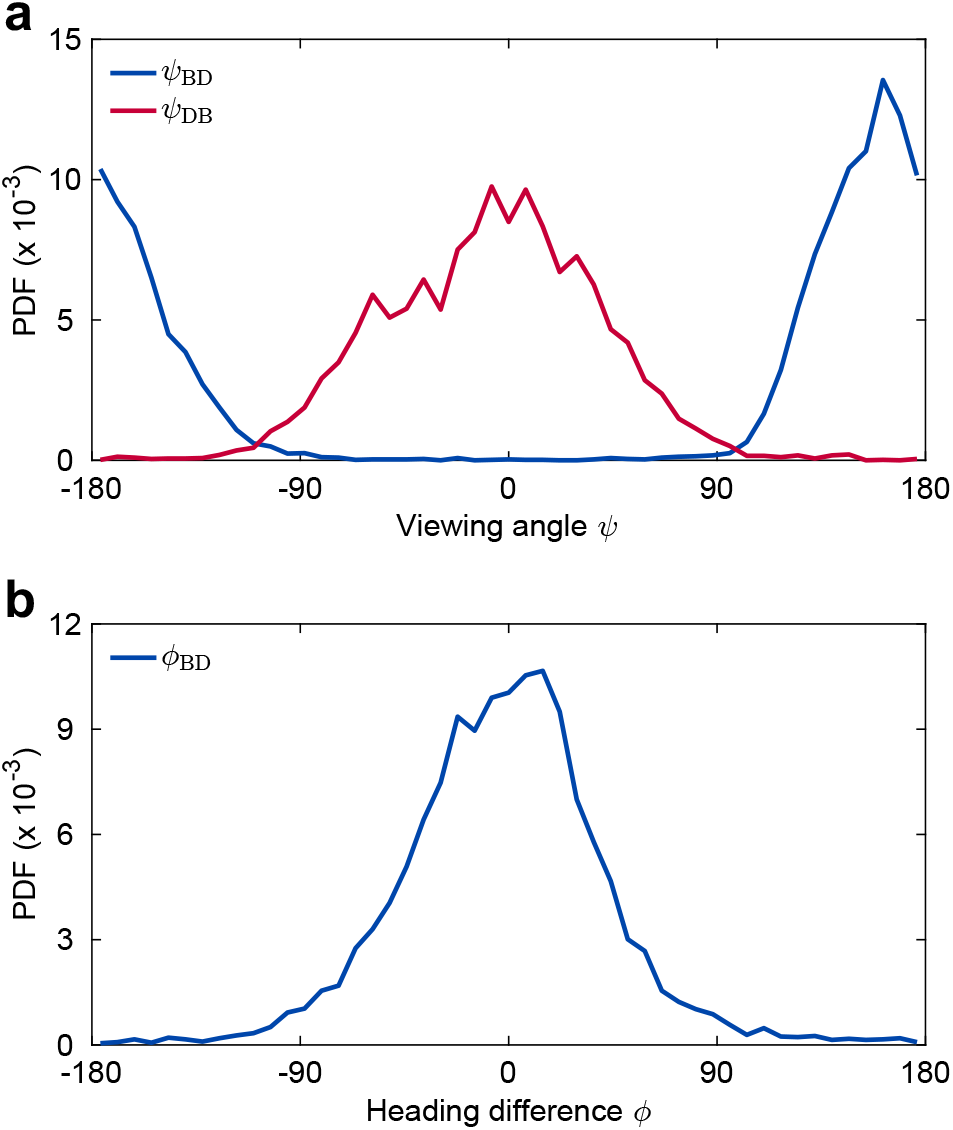
Probability density functions (PDF) of (**a**) the respective viewing angles *ψ*_BD_ and *ψ*_DB_ with which the barycenter (blue) and the dog (red) perceive each other, and (**b**) their heading angle difference ϕ_BD_ = ϕ_D_ − ϕ_B_ (blue).

We also calculate the turning rate, *δ*ϕ_*i*_ = ϕ_*i*_(*t* + Δ*t*) − ϕ(*t*), of the barycenter of the flock and that of the dog as a function of *ψ*_DB_ and *ψ*_BD_ respectively. The turning rate of barycenter of the flock (*δ*ϕ_B_) is close to 0 when the dog is behind the sheep (|*ψ*_BD_|≈ 180^∘^), but turns in the opposite direction of dog as |*ψ*_BD_| decreases, *i*.e. when the dog moves towards the flanks of the flock (Figure 5**a**).

**FIG. 5.**
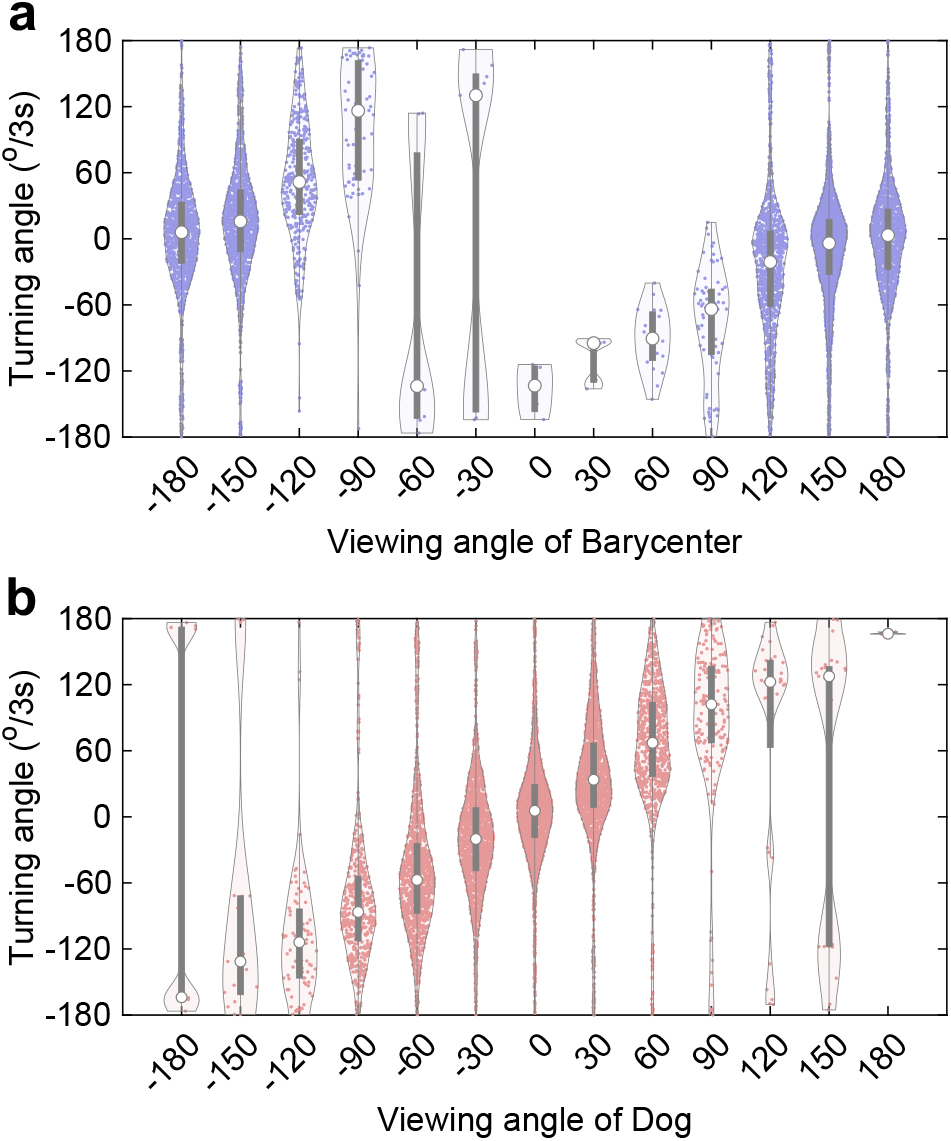
Turning angle over 3s of the barycenter of the flock (**a**) and the dog (**b**) with respect to their corresponding viewing angles *ψ*_BD_ and *ψ*_DB_. In all plots, purple dots represent the observed turning rates for a given viewing angle, white dots correspond to the median, and the thick vertical line corresponds to the limits of first and third quartiles respectively

Moreover, during herding events the dog turns towards the barycenter of the flock, and its turning rate (Δϕ_*D*_) is proportional to dog’s viewing angle (*ψ*_*DB*_); in response to orders given by the shepherd, the dog performs enveloping movements to the right or left, until sometimes its direction of movement is perpendicular to that of the flock (Fig. 5**b**). There is also a notable correlation between the speeds of the dog and the barycenter (Fig. 6**a**), and the flock becomes less cohesive as sheep move at higher speeds (Fig. 6). We find that both dog speed and barycenter speed increase or decrease simultaneously, suggesting a mutual influence between the dog and the sheep (Supplementary Fig.4). In the next section, we delve further into revealing the influence between the dog and the sheep at fine time scales.

**FIG. 6.**
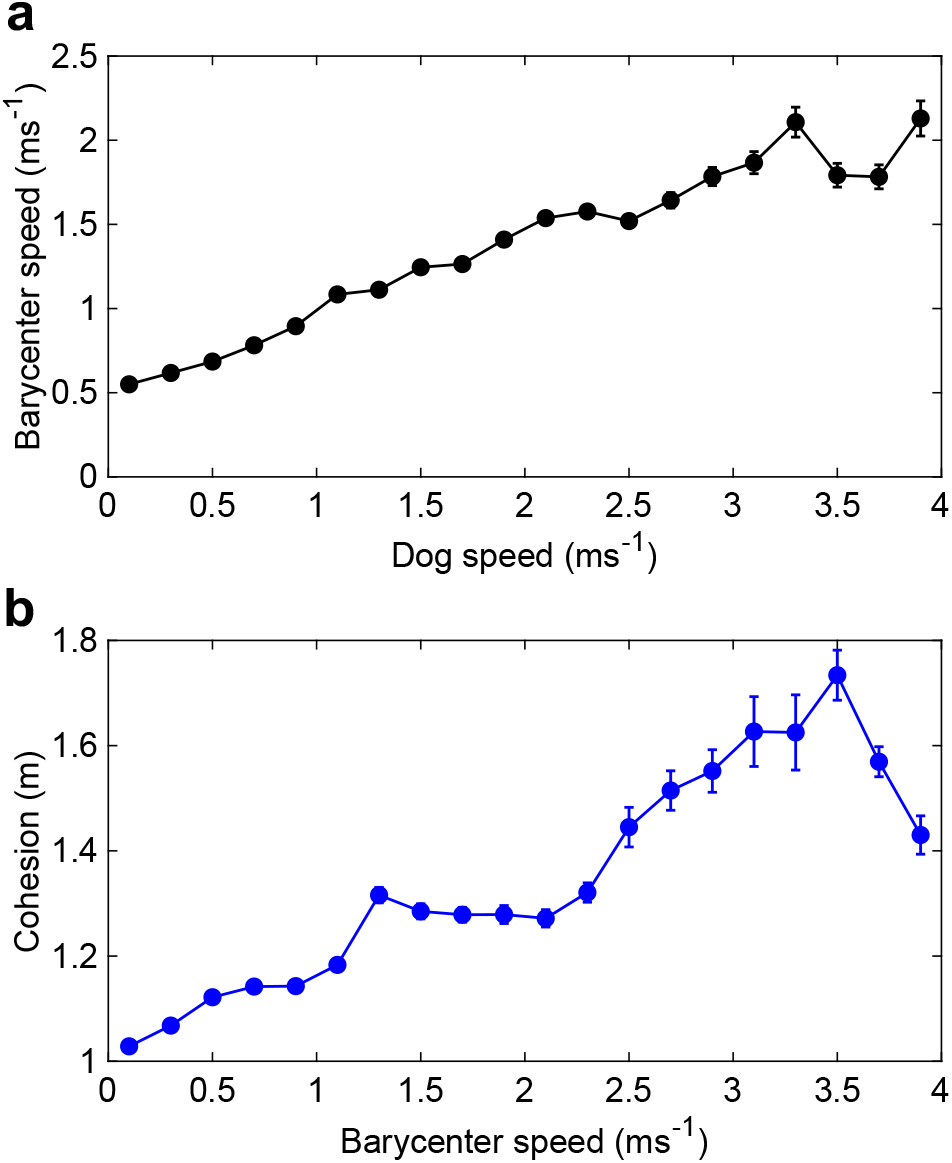
Speed of the barycenter of the flock as a function of the speed of the dog (**a**). Average group cohesion of the flock for varying group speed (**b**). As the sheep move faster, the flock becomes less cohesive. Error bars represent standard error.

### B. sDirectional correlations and hierarchical dynamics

In our experiments, the dog actively steers the flock between two designated points in the field. During these events, we aim to explore both the influence of dog behavior on the direction of sheep movement and how the behavior of individual sheep affects the direction of movement of its neighbors. We infer that an individual *j* follows *i* if *j* consistently copies *i*’s direction with some delay, denoted by τ_*ij*_. To identify this leader-follower relationship, we follow the methods described in [50–53] and calculate the cross-correlation function between orientations of *i* and *j*, henceforth referred to as directional correlation (Fig. 7a,b, Supplementary Section 1, Supplementary Fig.5).

**FIG. 7.**
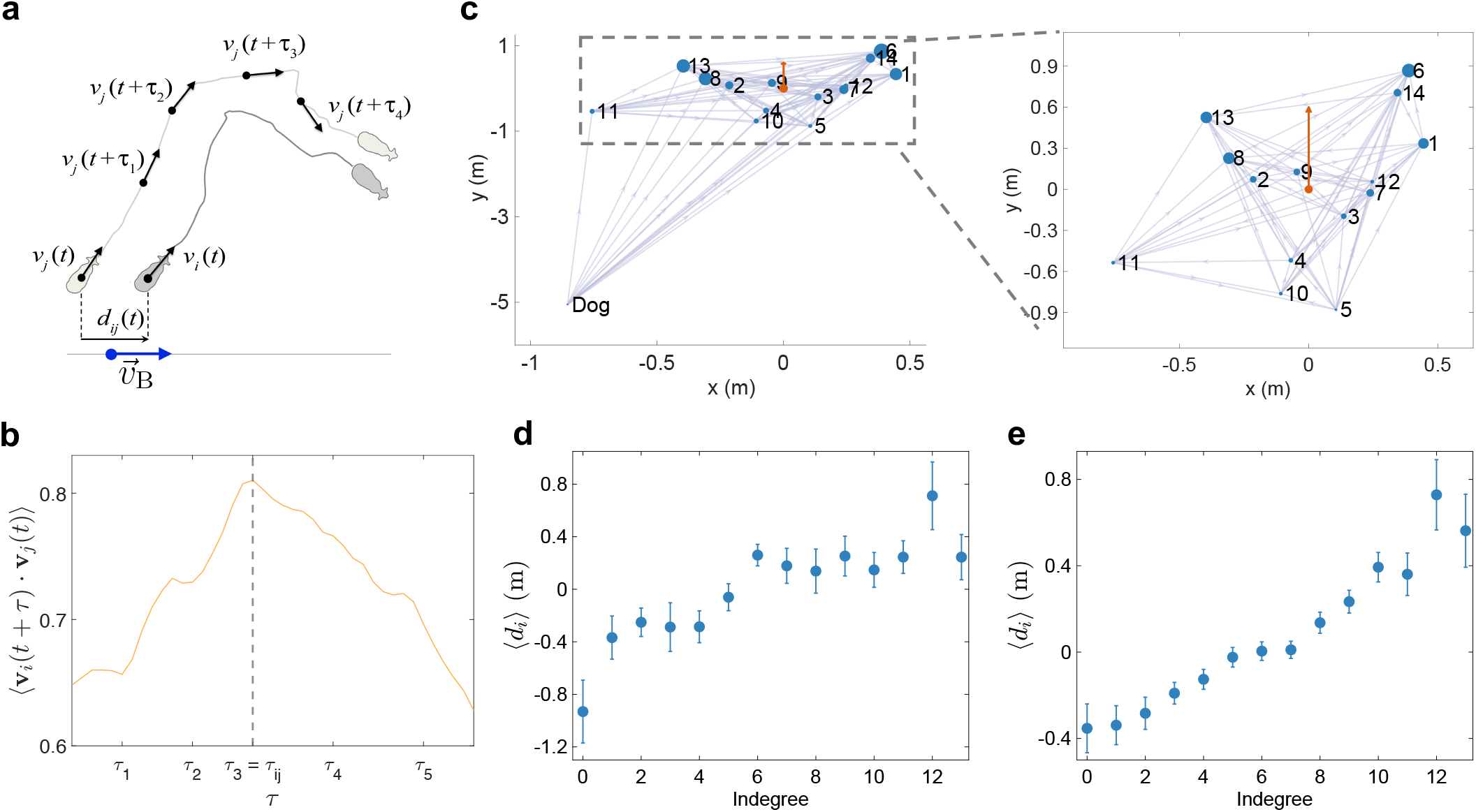
Graphical representation of normalized velocity correlation function analysis to identify leader-follower relationships. (**a**) *d*_*ij*_ is the projected relative distance 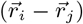 between individuals *i* and *j* onto the direction of motion of the sheep flock 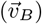 at time, t. (**b**) For each pair (between sheep, and sheep and dog), the velocity correlation is 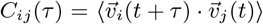 between 2 sheep during one of the chasing events. τ_*ij*_ is the correlation time delay, *i*.e. the time delay at which *C*_*ij*_ is maximum. (**c**) Leader-follower network of one of the drives. Nodes labeled from 1-14 are sheep and labels remain same throughout the experiment. Nodes are plotted at average distance from group center. Group center (orange dot) is at origin facing north. Edge is drawn from follower to leader and size of node is proportional to number of followers. Correlation between the average relative spatial position (mean *d*_*i*_ ± SE) and the hierarchical leadership network obtained from all chasing events in the data (**d**) and the model (**e**). Indegree serves as a proxy for hierarchy, where individuals with a high indegree are higher in the hierarchy.

We then construct a directed leader-follower network for each of the herding events based on the pairwise τ_*ij*_ values computed for that herding event. In such a network, each node represents either the dog or an individual sheep, and a directed edge is drawn from the follower to the leader. We infer the ‘leadership hierarchy’ of an individual by by computing the indegree (defined as number of incoming edges into a node) for the node representing the individual on the network. The higher the indegrees (or, in other words, number of followers) the higher the position in the leadership hierarchy.

Here, we note that, for a given pair, a leader-follower relationship may not always exist. Unless one individual in the pair consistently copies the direction of the other, the strength of directional correlation will be negligible. However in our data, we do observe consistent leader-follower pairs within each herding event. Figure 7c shows the leader-follower network observed during one of the herding events and Supplementary Fig.6. shows the networks observed during other herding events.

To understand leader-follower relationship further, we overlay the network on a reference frame whose origin is the centroid of sheep flock. Furthermore, in Fig.7c, the size of the node is proportional to its indegree and thus reflects its position in the leadership hierarchy. We observe that sheep in the front of the flock are characterized by a higher indegree (bigger node size) compared to the sheep located in the back, lastly followed by the dog. In other words, although the dog was driving the flock, it comes the lowest in the leadership hierarchy. This suggests that the dog continuously adjusts its movement direction based on sheep motion. Consequently, we expect that sheep in the front influences the group direction and, in turn, the direction of the dog.

To quantify the influence of individuals located at the front of the flock on the direction of its movements, we calculated the average distance of sheep from the group barycenter projected onto the group velocity, 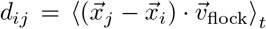, and 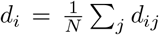, where *N* is the number of sheep (Fig. 7a). For all sheep that are in the front of the group, *d*_*i*_ > 0, and consequently, *d*_*i*_ < 0 for sheep at the back. As noted earlier, we use the indegree of a node as a proxy for its directional influence. So, we calculate *d*_*i*_ for all individuals with a given indegree. This informs us about the average spatial position of all sheep with the same indegree. In simple terms, it indicates where the sheep is located (front or back of the group center) for a given indegree. We find a strong correlation between *d*_*i*_ and the indegree of a node confirming that sheep located in the front of the flock influence the direction of motion of the group, and in turn, on dog movement (Fig.7d, Pearson correlation for indegree versus ⟨*d*_*i*_ ⟩, ρ = 0.85, P = 0.00004). In addition, Supplementary Fig.7 shows that some individiuals are consistently found at the front of the group during the different herding events.

### C. Modeling the collective response of the flock to the herding dog

To better understand how information propagates in the flock as a consequence of social interaction between sheep and the interactions with the dog, we use a discrete time model where the positions of individuals are given at equispaced time instants *t*^*n*^ = *n*Δ*t*, Δ*t* = 1 s. This herding model is adapted from a model developed by Strömbom et al [54].

#### Sheep

At a given time *t*^*n*^, each individual sheep *i* is located at 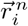 and moves to the point 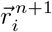 given by

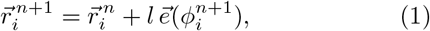

where 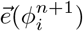 is the unit vector in the direction of 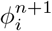 and l is the length traveled during this period of time. The angle 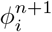 is the sheep heading’s angle during this *n*th time step and is given by the direction of the weighted additive combination of the vector director of the previous time step 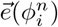, an additive noise 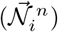, and the vectors corresponding to the external interactions to which the sheep is subject,

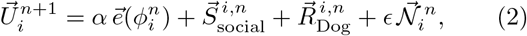

where 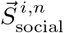 are the social interactions between sheep, 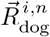 the repulsion from the dog. We assume that interactions to which individuals are subject can be expressed by linear functions that do not depend on the distance separating them.

Usually, in self-propelled particle models, agents align and attract with *all* the nearest, topological or metric neighbors [55]. However, recent studies have shown that in a moving animal group, each individual does not interact with all its neighbors but only with a small number of them [56–58]. This selective attention reduces the amount of information that needs to be processed by an individual thus avoiding cognitive overload. For instance it has been shown that fish only interact with their most or two most influential neighbors [49, 59, 60] or one randomly chosen neighbor [61]. It has also been shown that sheep select interaction neighbors [62]. To incorporate this cognitive constraint in our model, we introduce a key modification — sheep perceives a limited number of its nearest neighbors, among which only *k* have an impact on its behavior [63].

Thus, a number *n*_Att_ of these neighbors, chosen randomly, are considered to attract the sheep, and to contribute equally to the strength of the attraction 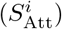 and a number *n*_Ali_ ≤ *n*_Att_, and sampled randomly from the *n*_Att_ attracting ones, are considered to act on the alignment of the sheep 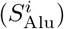, all of them with the same intensity. Finally, the sheep is repulsed by every sheep closer than a short distance *d*_Rep_, and this, with the same intensity. Thus, social interactions between sheep are described as follows:

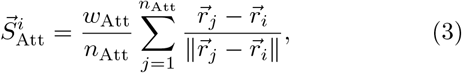

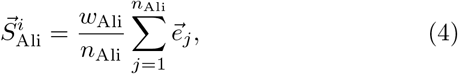

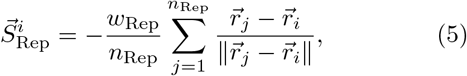

where each interaction has a positive weight *w*.

When the dog is closer than a distance *R*_*D*_, the sheep is repulsed in the opposite direction according to

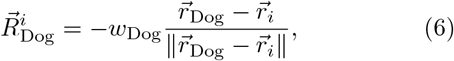

where *w*_dog_ is a positive weight.

#### Dog

Sheepdogs are typically controlled by a shepherd, who wishes to drive the flock from one point to another by using the dog as a repulsive stimulus that makes the individual sheep to move away from it. We are interested in situations where the sheep are grouped in a flock and the sheepdog is positioned at one side of the flock. Then, the shepherd adjusts the position of the dog to make the flock travel towards a target point *T* . For simplicity, we assume that the dog moves straight with constant speed *v*_D_ = 1.5 ms^−1^, unless a sheep is closer than a short distance l_*a*_, in whose case the dog slows down to *v*_D_ = 0.05 ms^−1^.

At each time step, dog’s position is updated as a func tion of the position of the barycenter of the flock with respect to the target point, so that the repulsion that dog exerts on the sheep points towards the target. Thus, the dog moves towards a point *P*_drive_ such that barycenter of flock B lies between *P*_drive_ and the target *T*, ensuring that the three points are aligned and that a distance l_drive_ separates *P*_drive_ from B (Fig. 8). In case that an individual sheep separates from the flock more than a distance l_sep_, the dog has to make it to come back to the group. Then, *P*_drive_ is defined in the same way but considering that the barycenter of the flock is the target and *S*_*i*_ is the sheep to be driven. Once the sheep is back to the group, the dog returns to drive the flock towards the target.

**FIG. 8.**
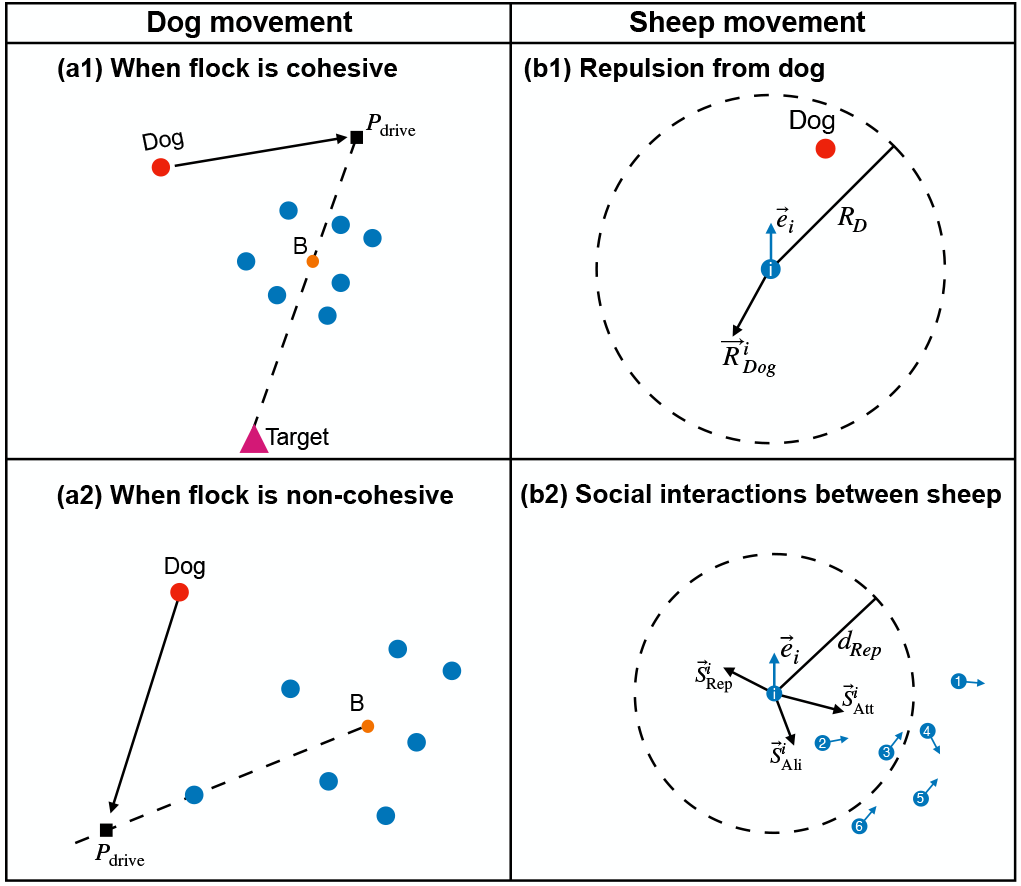
Schematic representation of the interaction rules in the computational model for (**a**) dog and (**b**) sheep. At each time step, the dog’s position is updated based on whether: **a1)** the flock is cohesive (*i*.e. the distance of the farthest sheep from the barycenter (B) is less than l_*sep*_), or **a2)** non-cohesive (*i*.e. the distance of the farthest sheep from the barycenter is greater than l_*sep*_). The dog moves in a straight line at a constant speed, *v*_D_, if it is farther than *d*_Rep_ distance from all agents. Otherwise, it moves at 0.0075*v*_D_. (**b1**) Sheep move away (along 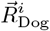) from the dog if it is within distance *R*_D_. Sheep are re from their neighbors within a distance of *d*_Rep_ Along 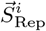. Sheep align and attract towards a set of randomly chosen neighbors among those they can perceive. For example in (**b2**), the focal sheep can perceive five nearest neighbors (*k* = 5, marked 1-5). The focal sheep is attracted towards the average direction, 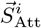, of four randomly chosen neighbors (*n*_Att_ = 4, sheep marked 1-4 in this case) and aligns (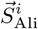) with one of those chosen neighbor (*n*_Ali_ = 1, sheep marked 4 in this case). We set *k* = 10, *n*_Att_ = 5, and *n*_Ali_ = 1 in our simulations (see Section.III C). The resulting heading vector of sheep *i* is a linear weighted combination of repulsion from the dog, collision avoidance, attraction towards neighbors, orientation towards neighbors, each with their corresponding weights given by *w*_Dog_, *w*_Rep_, *w*_Att_, and *w*_Ali_ respectively. We set *R*_*d*_ = 12, *R*_*a*_ = 2, *w*_Dog_ = 1, *w*_Rep_ = 2, *w*_Att_ = 1.5, *w*_Ali_ = 1.3, α = 0.5, e = 0.5, *v*_S_ = 1 ms^−1^, and *v*_Dog_ = 1.5 ms^−1^ in our simulations.

The point *P*_drive_ is then given by

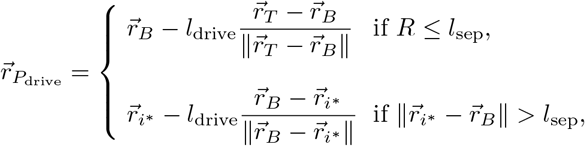

where *i*^*\**^ is the most distant sheep from the barycenter, 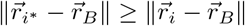 for all *i* = 1, …, *N*.

Supplementary Fig.8 shows the results of extensive numerical simulations of the model (see also Supplementary Videos 3 and 4) and the values of the parameters used in the model are listed in Supplementary Table 1. We find that group properties such as polarization, cohesion, and elongation are in qualitative agreement with experimental data. Additionally, we find that the PDFs of viewing angle of dog and barycenter, and the difference in heading angles between group barycenter and the shepherd qualitatively agrees with experiments. We also construct the interaction network as described in Section III B. Here too we find that individuals in the front of the flock are more often influential in directional decisions, with the dog consistently adjusting its direction in agreement with experimental observations (Fig. 7e).

Furthermore we aim to understand if the observed pattern of information propagation from front to back results from a constant perturbation induced by the dog or if it merely emerges as a consequence of social interaction between sheep. To study the precise influence of the dog, we construct a null model of collective movement wherein we exclude the dog from the model described above. In the null model, sheep follow the same behavioral rules as in the herding model. However sheep do not graze but always move at a constant speed following a combination of local interaction rules: moving towards near neighbors, aligning with their average direction of motion and avoiding collisions. In that case, when we construct the interaction network based on directional correlations, we do not find any leader-follower pairs. Even when they exist, there are only 2-3 pairs of leader-follower and the leader-follower network is not as highly connected as the one observed in the real data or the herding model (see Supplementary Fig.9). As there is no inherent hierarchy between the individuals in the model, and as they do not occupy a fixed spatial position within the flock during unperturbed collective movements, we would expect that there would be no consistent leader-follower pairs. However, in the presence of a constant perturbation such as a herding dog, individuals cannot change their spatial positions frequently, resulting in a hierarchical transfer of directional information of group motion. Therefore, the individuals which are located at the front of the flock have more influence on its direction of motion only when they are consistently herded by the dog.

Lastly, it is important to notice that the last position occupied by the dog in the hierarchy with respect to sheep is not explicitly coded in the model but emerges as a result of the interactions between the dog and sheep. This confirms that simple interaction rules of collecting and driving can reproduce the empirically observed patterns, at least in small herds.

## IV. DISCUSSION

We investigated the collective responses of a flock of merino sheep interacting with a herding dog. Specifically, we study (1) the influence of the dog on the direction of sheep movement and (2) the impact of individual sheep both on their neighbors and the dog. By an analysis of high-resolution spatiotemporal tracks of both the dog (acting as a threat or stressor) and the sheep flock, we characterize the collective response behavior of sheep as well as offer novel insights on the hierarchical nature of directional information flow.

In our experiments, we observe a highly polarized and cohesive sheep flock throughout the herding events. Additionally, we observe alignment between the flock and the dog. We emphasize that the groups are not only highly cohesive but also exhibit high polarization. Moreover, as the dog increases its speed, the group increases its own speed while being highly polarized but less cohesive. The increase in group cohesion observed in our experiments when sheep interact with a herding dog is consistent with the findings of King et al [43]. However, recent studies indicate that the selfish herd alone may not adequately explain the anti-predatory benefits of group living, especially when groups exhibit synchronous collective motion as observed in our sheep flock experiments [9, 18, 41]. Furthermore, computational models suggest that when the speeds of predators and prey are comparable, individuals aligning with neighbors rather than moving towards the group center are less likely to be captured [64].

Through the analysis of time delays of directional correlations, we identify a clear hierarchy among sheep in terms of their directional influence on the flock. We find that the average spatial position of a sheep along the front-back axis of group velocity is strongly correlated with its influence on group movement. In other words, although the flock is continuously chased by the dog, we find that the dog aligns its movement direction, on short time scales (∼ seconds), with that of the flock, with the directional information flowing from front of the flock to the rear and then to the dog. While this appears counter-intuitive, in a previous study, Early et al. [65] speculated that the dog’s adjustment was based on the flock’s movement in a yard. This unexpected directional hierarchy likely results from the close proximity between the dog and the flock in confined sheep yards, which allows the flock to move in its intended direction while the dog adapts to the herd on a short time scale.

We also observe that some sheep find themselves typically in the front of the group. We suggest that this spatial heterogeneity in sheep positions within the flock may arise from differences in individual reactivity to dogs [66], with the individuals that are most sensitive to the presence of the dog are those who are at the front of the flock, thus initiating the directional changes.

While it is clear that individuals located in front had a greater directional influence on the herd, we did not observe a significant correlation between the spatial position of an individual and its influence on the speed changes of neighboring sheep (Supplementary Fig.10). This could be explained by the fact that as the flock maintains its cohesion during the herding events, it imperative that individuals move at a similar speed. Therefore, there may be no preferred position within the flock to influence the group’s speed or that of any other flock member. However, as shown in [35, 67], the directional changes in escape events do appear to depend on the spatial location of individuals since those on the periphery have more degrees of freedom to alter the direction of motion.

Our findings are consistent with observations in mobbing flocks where individuals influencing the group’s direction of motion were located at the front [51], however the study used a fixed ground-based predator model. Similar front-to-back directional information flows have also been reported in homing pigeons and meerkats exhibiting directed collective motion, although individuals were not subjected to an external threat [50, 68]. On the other hand, in jackdaw flocks exhibiting synchronized motion but without any perturbation, the birds influencing the group’s direction were not only found at the front or edges but also at the rear [51]. In meerkat groups in motion, the individuals located in front do not influence the speed of the group [68]. All this suggests that it is essential to consider the ecological context when inferring the social or hierarchical influence of individuals on each other in animal groups.

We strengthen our inferences with the use of a computational model to show that sheep following simple interaction rules (*i*.e. repulsion from the dog and a tendency to move towards nearby neighbors and align with them), can reproduce collective response patterns similar to those observed in our experiments. Specifically, we observed a correlation between the average spatial position of sheep within the group and their hierarchy in directional influence. While certain individuals are often observed at the front of the group in experiments, the model suggests that it is not necessary that individuals have fixed spatial positions within the flock to obtain the experimentally observed pattern of correlation between the spatial position of sheep and its hierarchical influence in the movement direction. In addition, using a null model, we also show the presence of a constant external perturbation that drives away the flock, such as a herding dog, is a must to produce the hierarchical directional influence from front to back.

Our observations also reveal that the flock is often elongated perpendicular to the group velocity. This finding contrasts with expectations from fish schools, where oblong formations (*i*.e. the group is elongated along its direction of motion) are considered to provide protection against predation [69]. The analysis of the herding model shows that group elongation is determined by the relative strengths of attraction (*w*_Att_) and orientation (*w*_Ali_). The model predicts that if the orientation strength is lower than the attraction strength, the mode of elongation occurs at values less than 1 (Supplementary Fig.11); indicating that in sheep flocks, the strength of attraction is relatively higher than that of alignment. However, we emphasize that the alignment strength remains large to maintain a high degree of polarization observed in data, suggesting that both cohesion and highly directed motion are crucial for predator avoidance.

A logical continuation of this work would be to reconstruct the interaction rules between sheep, and between sheep and the dog directly from the UWB data. Interaction rules between individuals have been directly characterized in various species of schooling fish [29, 70]. Using a similar method and considering that Ubisense tags are both user-friendly and cost-effective, one can characterize and quantify interaction rules within this system as well. Studies on collective escape patterns suggest that fish schools may exhibit either subcritical or critical behavior [71–74]; a thorough characterization of interaction rules and phase-diagram of the underlying model can elucidate whether sheep flocks too exhibit features consistent with systems near criticality [75]. Such insights will enable the development of enhanced herding models with applications spanning various fields, including the creation of robots designed for environmental cleaning, crowd management, guiding groups of exploratory robots and bio-herding [76, 77].

## ACKNOWLEDGMENTS

We thank Gérard Latil for his help in preparing and running the experiments and Aitziber Ibañez for her help in the experimental design. We are deeply indebted to Ramón Escobedo for his participation to the experiments and for his assistance in pre-processing and analyzing the UWB data, and preparing the figures. We gratefully thank Mathias Aletru and his dog “donut” for their participation to this study. GT also gratefully acknowledges the Indian Institute of Science to serve as Infosys visiting professor at the Centre for Ecological Sciences in Bengaluru.

## Code and data availability

The codes for all the analyses and the computational model are available on the GitHub repository: https://github.com/teelab/collective-responses-of-flocking-sheep-to-herdingdog.git.

## Author contributions

GT designed the experiments. RP, CZ, MR, RB and GT carried out the experiments. VJ designed the model and performed numerical simulations. VJ, VG and GT analyzed the data and wrote the paper. All authors participated in discussing and approving the final version of the paper.

## Competing

The authors declare no competing interests.

