## Supplementary information for "Collective responses of flocking sheep to a herding dog"

#### Supporting information

- **Supplementary Figure 1-11**
- **Supplementary Table 1**
- **Legends for Supplementary Videos**

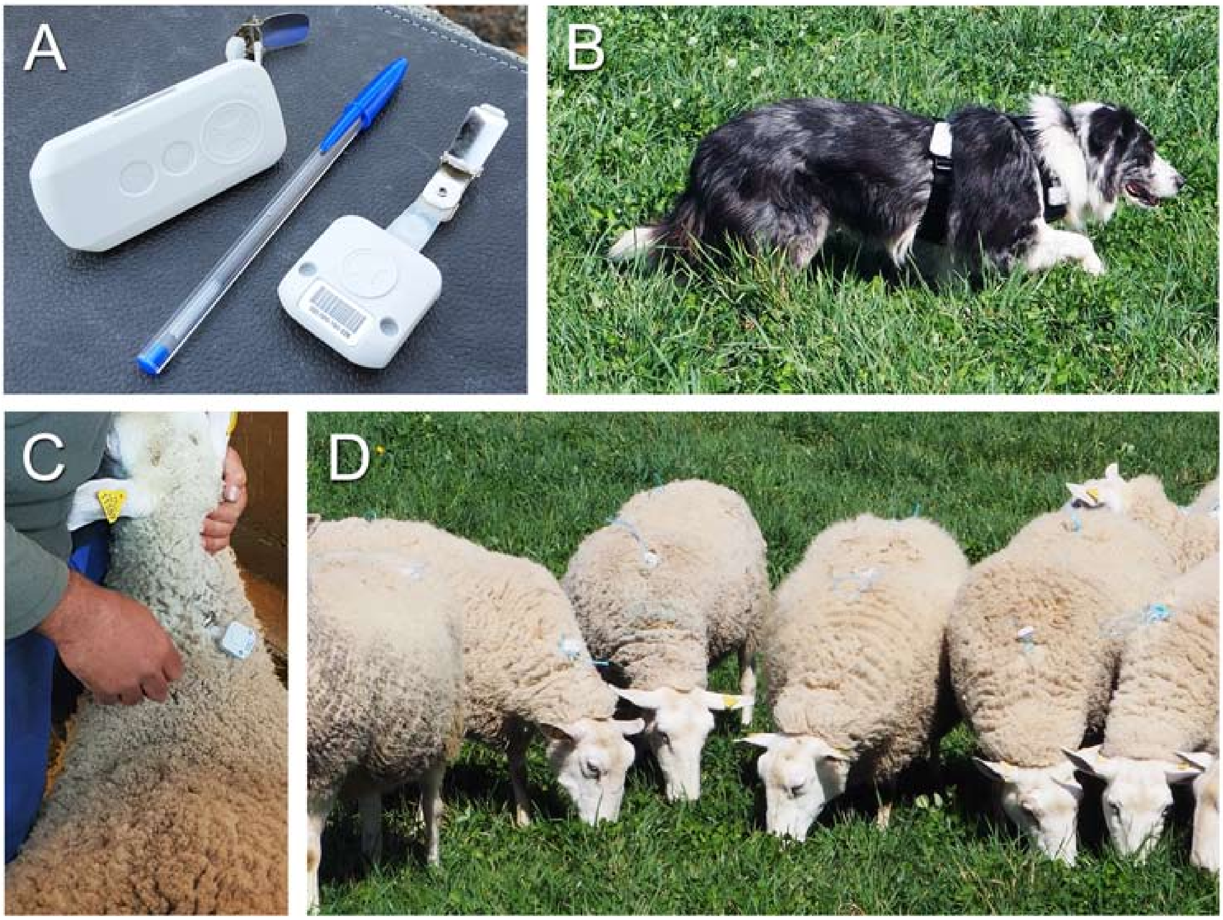


**Supplementary Figure 1.** (A) Ubisense tags used for tracking and tag attachment to the shepherd dog (B) and the sheep (C and D).

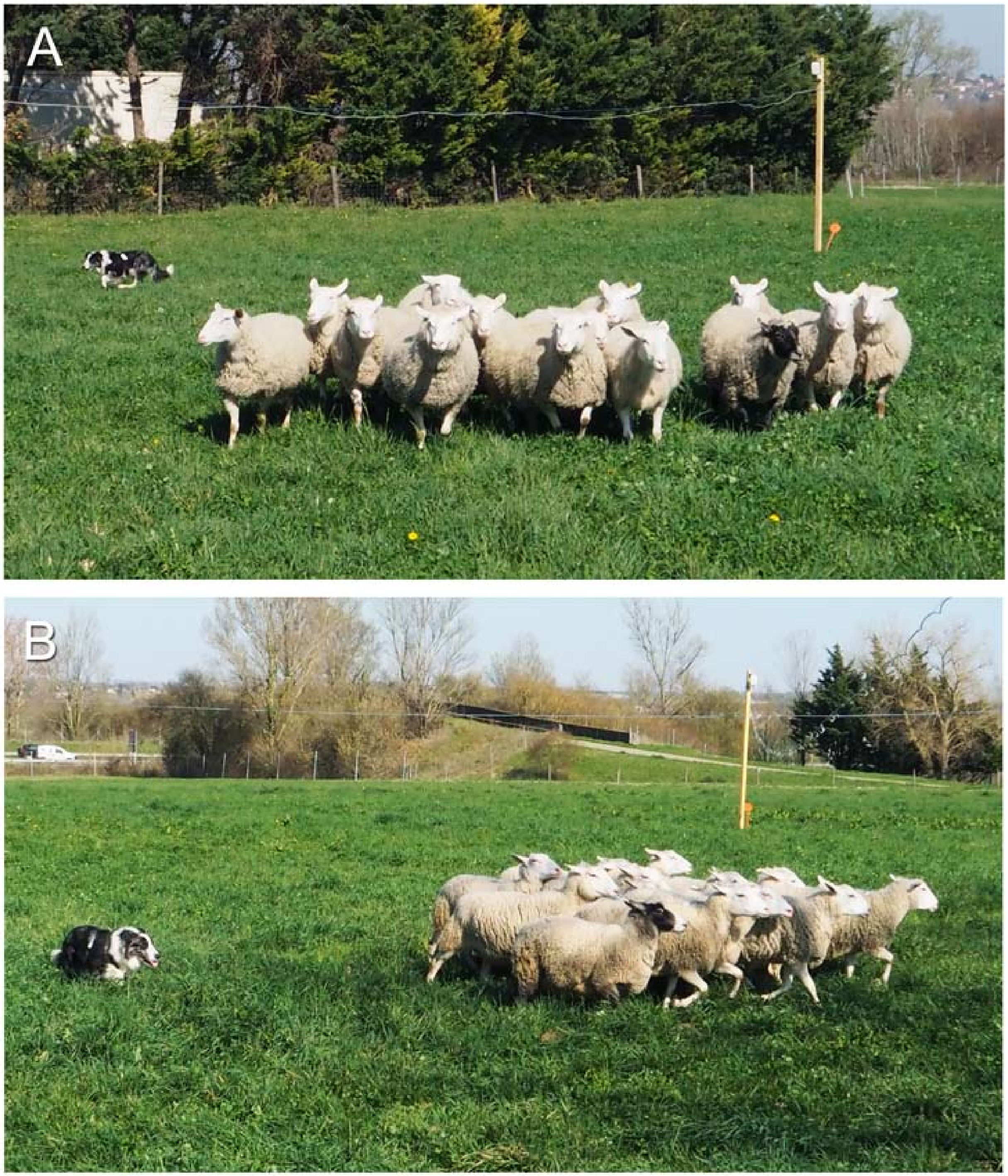


**Supplementary Figure 2.** Spatial formations adopted by the flock of sheep when it is chased by the dog with a group elongation $E < 1$ (A) and $E > 1$ (B).

### 1 Directional correlation function

The cross-correlation function between orientations of $i$ and $j$, referred to as directional correlation in the main text, for a time delay of $\tau$ is defined as $C_{ij}(\tau) = \langle \hat{v}_i(t+\tau) \cdot \hat{v}_j(t) \rangle_t$, where $\hat{v}_i$ is the orientation of $i$

and $\langle .. \rangle_t$ represents time average. The directional time delay, $\tau_{ij}$, is the delay at which $C(\tau)$ is maximum. Individual $i$ is considered a leader if $\tau_{ij} < 0$ or a follower if $\tau_{ij} > 0$. We consider that a pair $i$ and $j$ exhibit a leader-follower relationship only if $C_{ij}(\tau_{ij}) > C_{min}$ where we set $C_{min} = 0.5$.

#### 2 Cross-correlation between median speed of sheep, median speed of dog, and group polarisation

We evaluate the temporal cross-correlation between the following three scalar quantities: median speed of sheep, median speed of dog, and group polarisation. Cross-correlation between time-series of 2 scalar quantities $x_t$ and $y_t$, for a given time lag, $\tau$ is defined as,

$$C(x, y; \tau) = \frac{\sum_{t=0}^{t=T-\tau-1} (x_{t+\tau} - \bar{x}) (y_t - \bar{y})}{\sqrt{\sum_{t=0}^{t=T} (x_t - \bar{x})^2} \sqrt{\sum_{t=0}^{t=T} (y_t - \bar{y})^2}}, \quad (1)$$

where $T$ is the length of the time series, and $\bar{x}$ and $\bar{y}$ are means of time series $x_t$ and $y_t$.

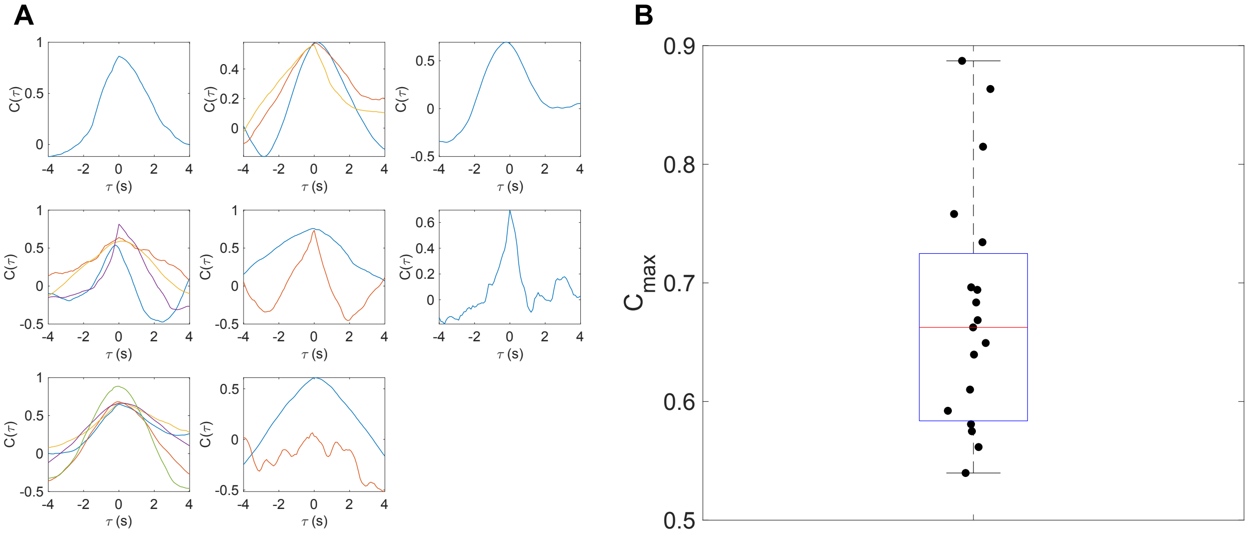


**Supplementary Figure 3. Cross-correlation between group polarisation and median speed of sheep**. (A) We calculate cross-correlation for 28 drives from 8 videos. Each line in a plot shows cross-correlations calculated for drives in the respective video. (B) Box plot of maximum value of $C(\tau)$ for each drive. We observe a positive correlation between group polarization and median sheep speed. In all box plots, red line correspond to median, and the blue horizontal line corresponds to limits of first and third quartiles respectively.

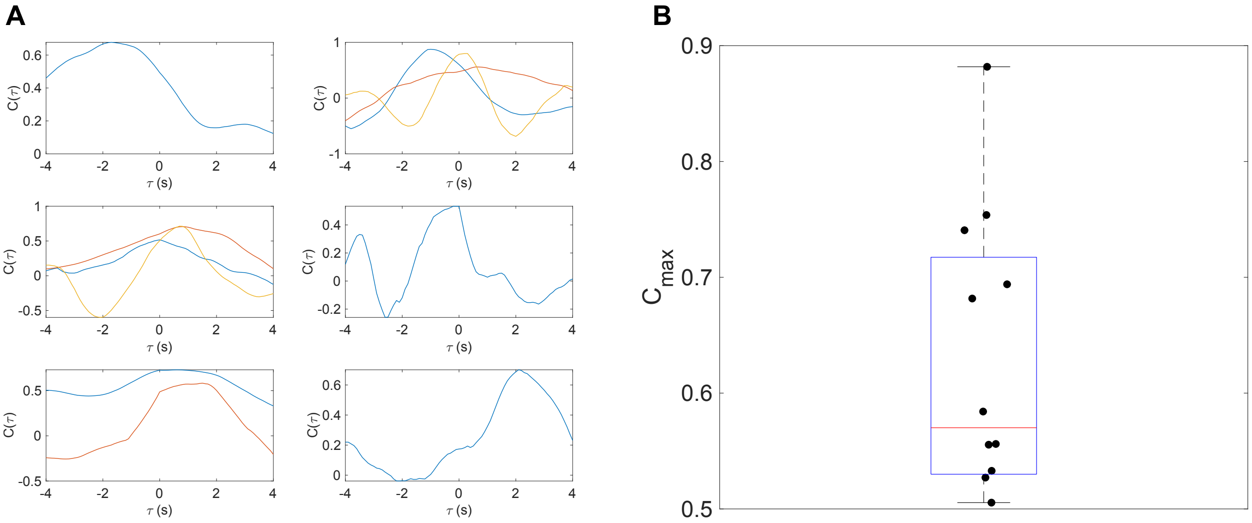


**Supplementary Figure 4. Cross-correlation between dog speed and median sheep speed.** (A) We asses cross-correlation for 28 herding events from 8 videos. Each plot represents cross-correlations for herding events in the respective video. (B) Box plot of maximum value of $C(\tau)$ for each drive. We observe positive correlation between dog speed and median speed of sheep. In all box plots, red line correspond to median, and the blue horizontal line corresponds to limits of first and third quartiles respectively.

### 3 Cross-correlation between normalised group velocity and normalised dog velocity

We define cross-correlation for time-series of 2 normalised vector quantities, $\vec{x}_t$ and $\vec{y}_t$, for time lag, $\tau$ as,

$$C(\vec{x}, \vec{y}; \tau) = \frac{1}{T - \tau} \sum_{t=0}^{T-\tau-1} \vec{x}(t+\tau) \cdot \vec{y}(t). \quad (2)$$

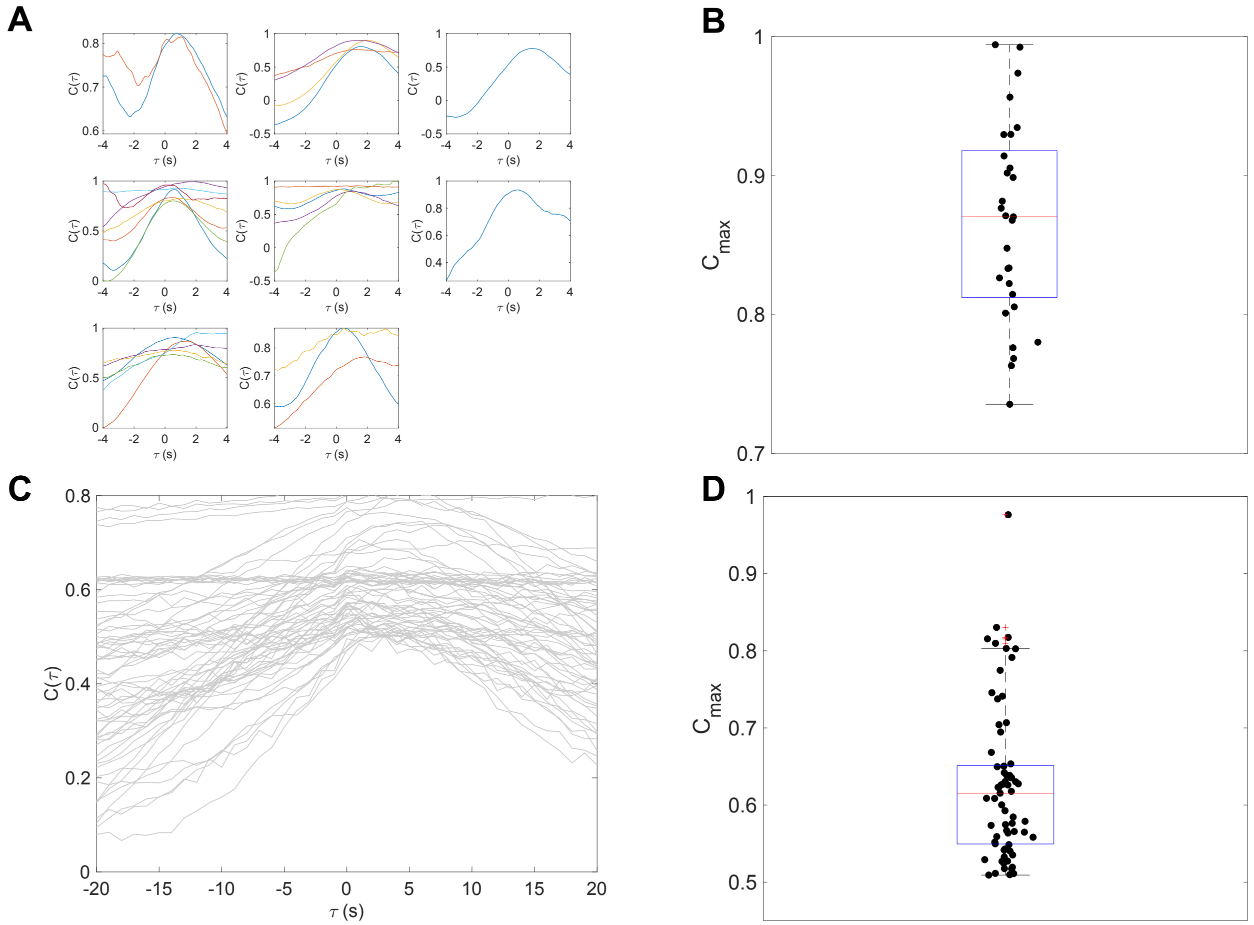


**Supplementary Figure 5. Cross-correlation between normalised dog velocity and normalised group velocity.** (A) Cross-correlation function for 28 herding events from all 8 videos. (B) Box plot of value of $C_{\max}$ for each drive. Normalized dog velocity and group velocity are positively correlated. We faithfully reproduce the results in the herding model. (C) Cross-correlation for 50 simulated chasing events. (D) Similar to empirical observations, we find a positive correlation between dog velocity and group velocity. In all box plots, red line correspond to median, and the blue horizontal line corresponds to limits of first and third quartiles respectively.

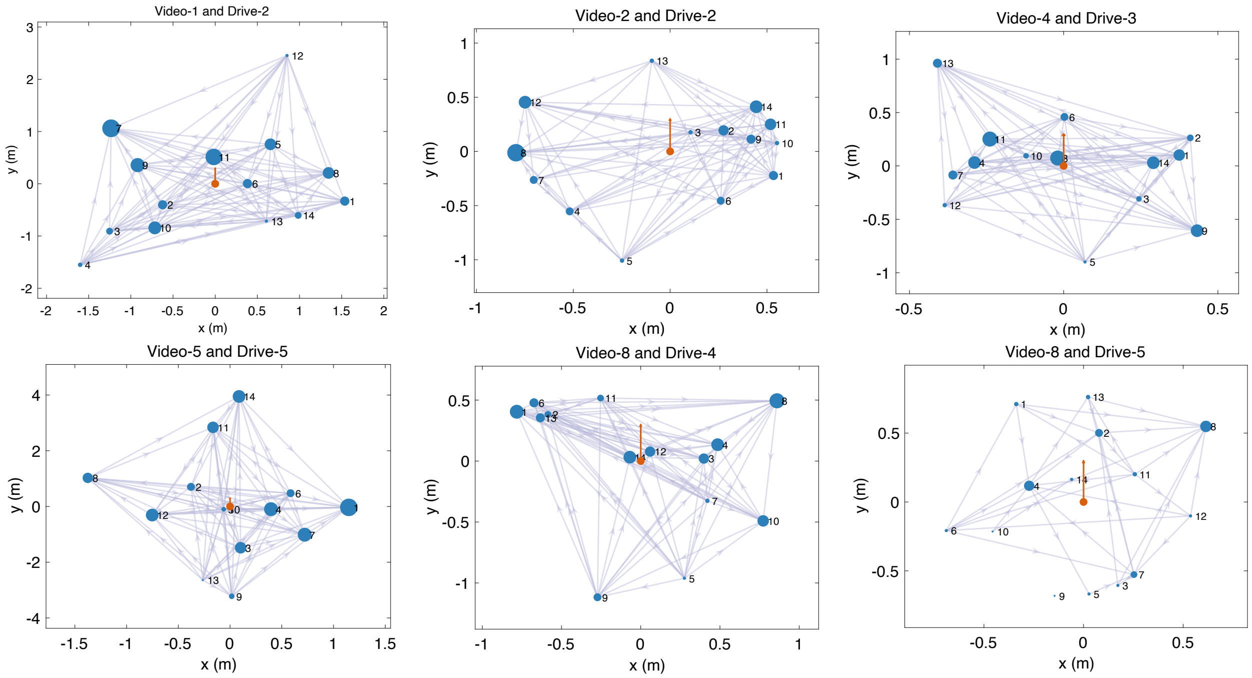


**Supplementary Figure 6. Leader-follower networks**. We construct a leader-follower network for all pairs of sheep based on velocity cross-correlations (see main text). Each graph corresponds to a herding event observed in a given video. Nodes labeled from 1-14 represent sheep, and labels remain the same throughout the experiment. Nodes are plotted at the average distance from the group center. The group center (orange dot) is at the origin facing north. An edge is drawn from the follower to the leader, and the size of the node is proportional to the number of followers. We observe that individuals in the front have a larger node size, indicating a clear hierarchy among sheep in terms of directional influence, with individuals in the front being more influential. We observe a clear correlation between spatial position of a sheep and its directional influence on the flock.

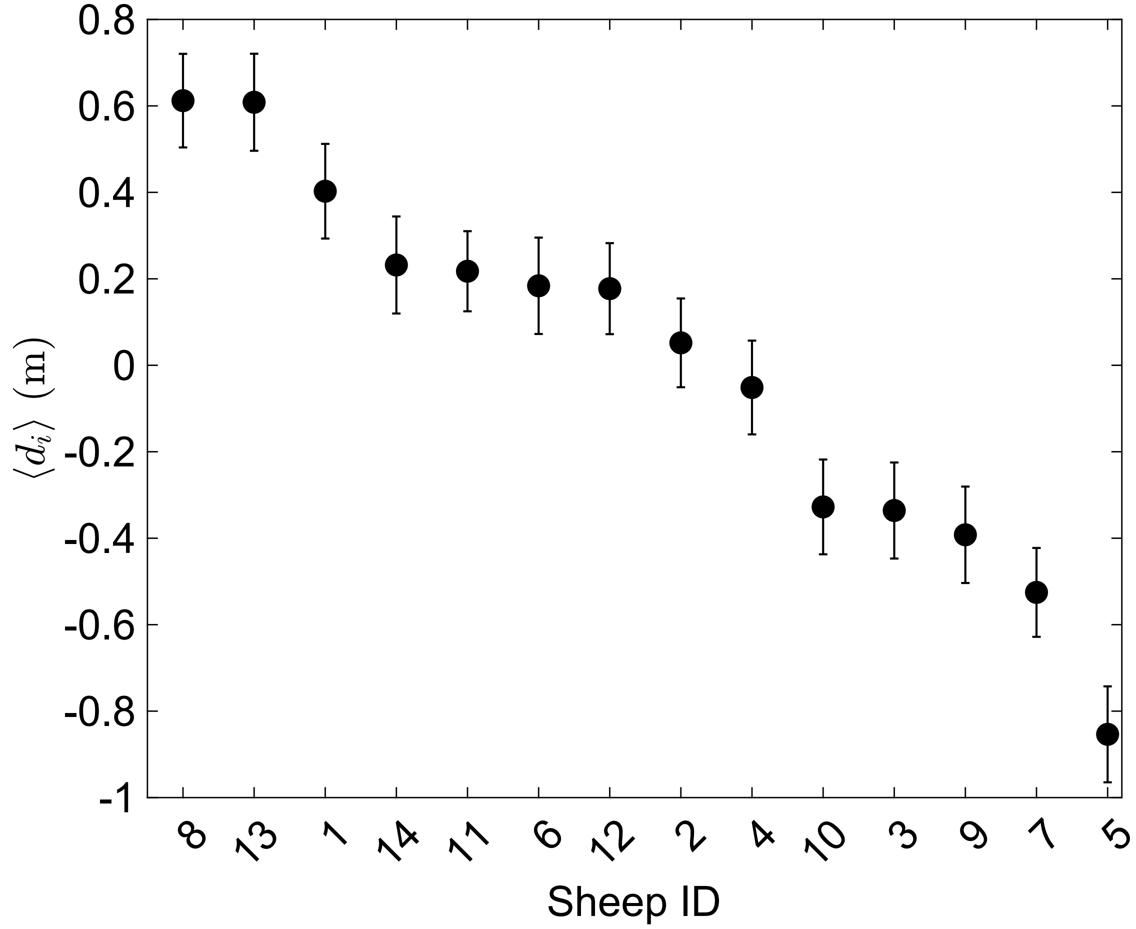


**Supplementary Figure 7.** Average relative spatial position (mean $d_i \pm$ SE) of sheep, ID from 1-14. We observe that some sheep are likely to be in the front (i.e., $d_i > 0$, sheep ID - 1, 8 and 13) while others are often found at the rare (i.e., $d_i < 0$, sheep ID - 5,7 and 9).

### 4 Model Results

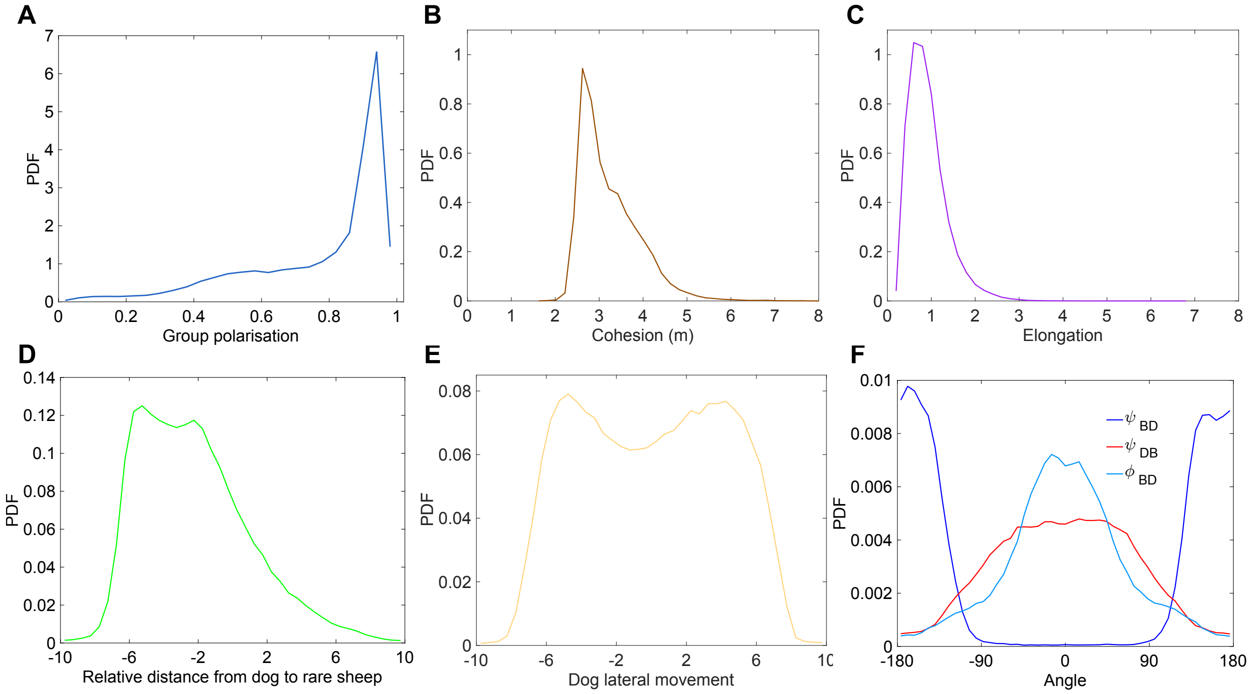


**Supplementary Figure 8.** Probability density functions (**a-f**) of the observables characterizing individual and collective behavior of sheep and their reaction to the dog in the simulations of the model. **a** Polarisation. **b** Cohesion. **c** Elongation. **d** Relative distance of dog to rear sheep. **e** Dog lateral movement. **f** viewing angles $\psi_{\mathrm{BD}}$ and $\psi_{\mathrm{DB}}$ with which the barycenter and the dog perceive each other, and **b** their heading angle difference $\phi_{\mathrm{BD}} = \phi_{\mathrm{D}} - \phi_{\mathrm{B}}$. We observe that group properties are in qualitative agreement with experimental data.

### 5 Null model

In the null model, sheep follow the same behavioural rules as those used in the herding model and described in the main text. However, sheep do not graze but always move at a constant speed. Therefore, sheep moves following a combination of local interaction rules: a tendency to move towards near neighbours, aligning the direction of motion with them and avoiding collisions. This model is similar to classical models of collective motions widely familiar in the field of collective motion.

We study two versions of the null model. In null model 1, all parameters have the same values as in the herding model. In that case, we observe that the flock is cohesive but not polarised ($P \approx 0$). In the null model 2, all parameters have the same values as in the herding model but we change the number of interacting neighbours to $k = n_{\mathrm{Att}} = 10$ and $n_{\mathrm{Alg}} = 5$, to obtain a cohesive and polarised flock.

We then construct the interaction network from the simulation data based on velocity correlations as explained in the main text. In the null model 1, we do not find any leader-follower pairs. In the null model 2, we find a few leader-follower pairs. However, the leader-follower networks are not as highly connected as those observed in the real data or in the herding model. A representative interaction network from null model is shown in Supplementary Figure 10.

As there are no inherent hierarchy among the individuals in the model, we would expect that there would be no consistent pair of leader-follower particularly when the flock is disordered as neighbours of an individual keep changing all the time. However, in highly polarised flocks, at least among some pairs and for a short interval of time, there is a possibility that some neighbours does not change place in the flock. Therefore, in some simulations we observe a few pairs of leader-follower. However the network is not as highly connected as that observed in experimental data.

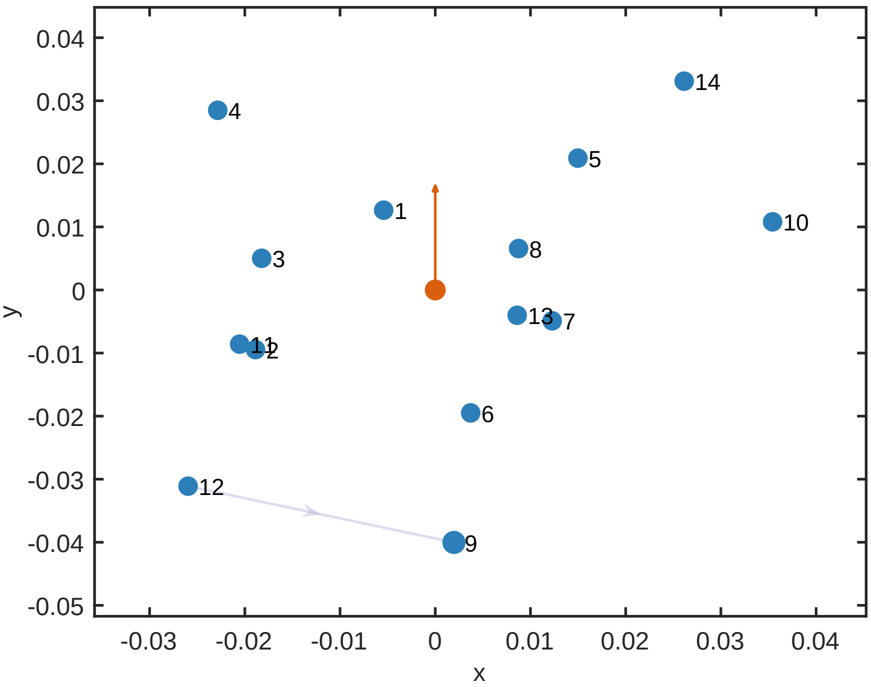


**Supplementary Figure 9.** Leader-follower network observed in null model. Nodes are plotted at average distance from group center. Group center (orange dot) is at origin facing north. Edge is drawn from follower to leader and size of node is proportional to number of followers. The leader-follower network is sparsely connected with only one edge from node 12 to node 9. The network is not as highly connected as observed in the real data or herding model.

### 6 Influence of spatial position on the hierarchical leadership network constructed based on speed cross-correlation

Here we access if there is a similar correlation between spatial position of a sheep and its influence on speed of the flock. First, we calculate the average distance of sheep from the group barycenter projected onto the group velocity as described in main text, $d_{ij} = \langle (\vec{x}_j - \vec{x}_i) \cdot \vec{v}_{\text{flock}} \rangle_t$, and $d_i = \frac{1}{N} \sum_j d_{ij}$, where $N$ is the number of sheep (see Figure 4A in main text). For all sheep that are in the front of the group, $d_i > 0$, and consequently, $d_i < 0$ for sheep at the back. Now we construct a leader-follower network for speed cross-correlations employing methods similar to velocity cross-correlations. We measure the time delay $\tau_{ij}$ where $C(s_i, s_j; \tau)$ is maximum. $s_i$ and $s_j$ are speed of sheep $i$ and $j$ respectively. We consider $\tau_{ij}$ for our analysis only if $C(\tau_{ij}) > C_{min}$. We set $C_{min}$ as 0.5. Individual $i$ is considered a leader if $\tau_{ij} < 0$ or a follower if $\tau_{ij} > 0$. From pairwise $\tau_{ij}$, we construct a directed leader-follower network for all chasing events. As described in main text, we calculate $d_i$ for all individuals with a given indegree evaluated based on speed correlations. Unlike for velocity correlations, we do not find any correlation between $d_i$ and the indegree of a node. (Figure.10, Pearson correlation for indegree versus $\langle d_i \rangle$, $\rho = 0.38, P = 0.24$).

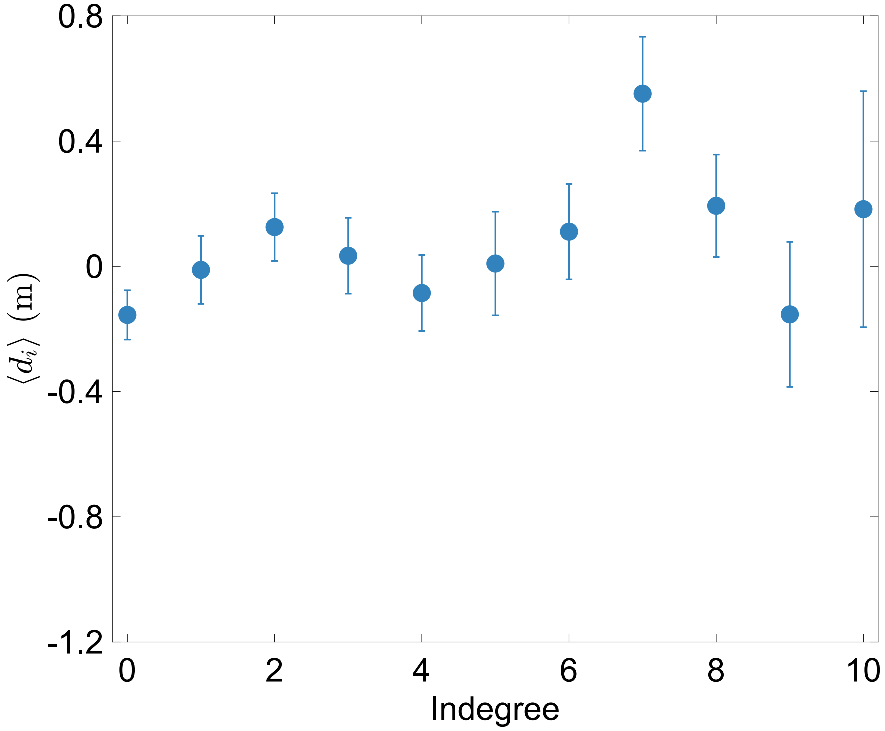


**Supplementary Figure 10.** Correlation between the average relative spatial position (mean $d_i \pm$ SE) and the hierarchical leadership network constructed based on speed cross-correlations obtained from all herding events in the data. The number of indegrees serves as a proxy for hierarchy, where individuals with high indegree are more influential. Unlike for velocity correlations, we do not find any correlation between $d_i$ and the indegree of a node.

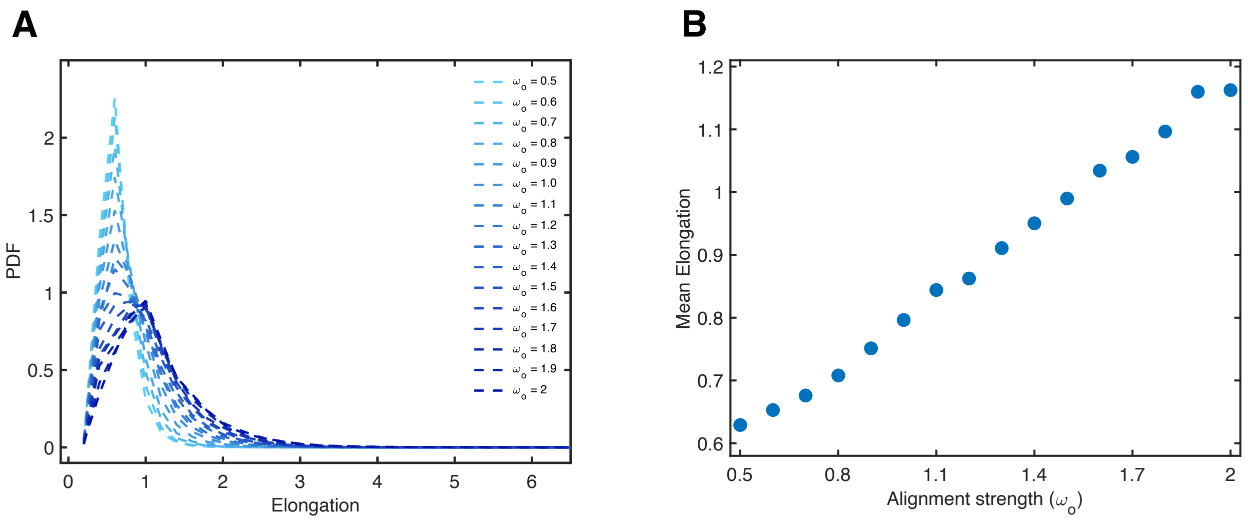


**Supplementary Figure 11.** **A**. Probability density function (PDF) of a group elongation for a range of alignment strength ($w_{\mathrm{Ali}}$) in the model. Other parameters are fixed to the values described in Table 1 of the main text. **B** For a fixed attraction strength ($w_{\mathrm{Atr}}$), we observe that group elongation increases with alignment strength ($w_{\mathrm{Alg}}$).

| Parameter | Description | Value |
|---|---|---|
| **Sheep** | | |
| $R_D$ | Maximum distance of dog detection (m) | 12 |
| $d_{\text{Rep}}$ | Between sheep repulsion distance (m) | 2 |
| $w_{\text{dog}}$ | Relative strength of repulsion from dog | 1 |
| $w_{\text{Rep}}$ | Relative strength of repulsion from other sheep | 2 |
| $w_{\text{Att}}$ | Relative strength of attraction towards other sheep | 1.5 |
| $w_{\text{Ali}}$ | Relative strength of alignment with other sheep | 1.3 |
| $k$ | Number of neighbors perceived by a sheep | 10 |
| $n_{\text{Att}}$ | Number of neighbors attracting the focal sheep | 4 |
| $n_{\text{Ali}}$ | Number of neighbors sheep randomly aligns with from $n_{\text{Att}}$ | 1 |
| $\alpha$ | Relative strength to move in the previous direction | 0.5 |
| $\epsilon$ | Relative strength of angular noise | 0.3 |
| $v_{\text{S}}$ | Sheep speed | $1\ ms^{-1}$ |
| **Dog** | | |
| $P_{\text{Drive}}$ | Driving position when flock is cohesive (m) | $l_d = r_d\sqrt{N}$ |
| $P_{\text{Drive}}$ | Driving position when flock is non-cohesive (m) | $l_c = r_d$ |
| $v_{\text{D}}$ | Dog speed ($\text{ms}^{-1}$) | 1.5 |
| $\epsilon$ | Relative strength of angular noise | 0.3 |

**Supplementary Table 1.** Model parameters. We chose model parameters such that flock remains cohesive as the dog steers it towards the origin. We fixed group size to $N = 14$ corresponding to the number of sheep in the experiments

### Supplementary videos

**Supplementary video 1:** Visualization of flock and sheep trajectories from a sample herding event, reproduced from UWB data. In the video, red, black, and green dots represent the dog, sheep, and shepherd, respectively.
**Supplementary video 2:** Visualization of flock and sheep trajectories from a sample herding event similar to Supplementary video 1. In the video, red, black, and green dots represent the dog, sheep, and shepherd, respectively.
**Supplementary video 3:** Numerical simulation of the herding model. In the video, red and black dots represent the dog and sheep, respectively. The flock consists of 14 sheep. The model parameters are detailed in Supplementary Table 1.
**Supplementary video 4:** Numerical simulation of the herding model. In the video, red and black dots represent the dog and sheep, respectively. The flock consists of 14 sheep. The model parameters are detailed in Supplementary Table 1.
